# Prolyl-hydroxylase inhibitor-induced regeneration of alveolar bone and soft tissue in a mouse model of ligature-induced periodontitis

**DOI:** 10.1101/2021.12.14.472623

**Authors:** Elan Zebrowitz, Tetsuhiro Kajikawa, Kamila Bedelbaeva, Azamat Aslanukov, Sam Bollinger, Yong Zhang, David Sarfatti, Jing Cheng, Phillip B. Messersmith, George Hajishengallis, Ellen Heber-Katz

## Abstract

Bone injuries and fractures reliably heal through a process of regeneration with restoration to original structure and function when the gap between adjacent sides of a fracture site is small. However, when there is significant volumetric loss of bone, bone regeneration usually does not occur.

In the present studies, we explore a particular case of volumetric bone loss in a mouse model of human periodontal disease (PD) in which alveolar bone surrounding teeth is permanently lost and not replaced. This model employs the placement a ligature around the upper second molar which leads to inflammation and bone breakdown and faithfully replicates the bacterially-induced inflammatory etiology of human PD to induce bone degeneration. After 10 days, the ligature is removed and the mice are treated with a timed-release formulation of a small molecule inhibitor of prolylhydroxylases (PHDi; 1,4-DPCA) previously shown to induce epimorphic regeneration of soft tissue in non-regenerating mice. This PHDi induces high expression of HIF-1α and the regenerative response is completely blocked by *siHIF1a*. Here, we observe that timed-release 1,4-DPCA rapidly and completely restores bone and soft tissue with normal anatomic fidelity and with increased stem cell markers due to stem cell migration into the site and/or de-differentiation of local tissue, PDL cell proliferation, and increased vascularization. In-vitro studies using gingival tissue show that 1,4-DPCA indeed induces de-differentiation and the expression of stem cell markers but does not exclude the role of migrating stem cells.

## Introduction

Humans, like virtually all mammals, heal tissue and organ injuries by the process of scarring with limited restoration of normal anatomical integrity and functionality. This is in contrast to species such as newts, salamanders and other vertebrates which heal perfectly through the process of regeneration. In these examples, bone and soft tissues are replaced creating indistinguishable replicas of lost or damaged tissues (1-3). There are several routes to mammalian regeneration being actively considered for regenerative therapies, usually involving the use of stem cells including those from the pulp as autologous grafts with or without bioscaffolds (4-9). There are also treatments with compounds such as high molecular weight hyaluronic acid which affect periodontal bone healing (10-12). However, the possibility of a systemically-acting drug which alone could induce regeneration in multiple tissues would be an intriguing therapy.

In the current set of studies, we explored the possibility that a prolyl hydroxylase inhibitor (PHDi), which regulates the expression of HIF-1*α*, can have a positive effect on the regeneration of bone and soft tissue of the jaw in mice. The use of the PHDi, 1,4-DPCA delivered in a hydrogel was previously shown by our laboratories to lead to a regenerative healing response resulting in the closure of ear holes and the replacement of soft tissue in mice. To study this PHDi’s effect on the jaw, we employed a mouse model of periodontal disease induced by placing a ligature around the upper second molar for 10 days, leading to an oral bacterial accumulation and bone degeneration, followed by removal of that ligature. 1,4-DPCA coupled to a PEG gel was then administered systemically followed by a second dose of drug administered 8 days later. After 20 days post ligature removal and the administration of drug, we saw full replacement of bone and soft tissue including the PDL which attaches the tooth to the bone.

The path to identifying such a drug began with the serendipitous observation that the MRL mouse strain, long employed in autoimmunity studies, possessed an unusual capacity for tissue regeneration. Through-and-through ear pinna wounds typically placed as life-long mouse identifiers healed with full closure and without scarring within 30 days. All tissue types found in the ear including epidermis, dermis, blood vessels, nerve, glands, cartilage and hair follicles were restored (13-14). Furthermore, multiple studies showed that this regenerative phenotype extended to MRL cornea, tendon, cartilage, muscle, fat, and other tissues (15-18). However, bone injuries were largely unexplored.

Insight into the biological basis of the MRL regenerative phenotype arose from the observation that the adult mouse employs a strongly aerobic glycolytic metabolism in the basal state similar to that seen in embryos. This metabolic state is enhanced during regenerative wound healing (19-20). One well-known molecule that regulates aerobic glycolysis is hypoxia-inducible factor (HIF-1α); this molecule is highly up-regulated in the MRL upon the initiation of injury, falling to pre-wounding levels after regenerative healing occurs (21). HIF-1α levels are regulated through the interaction with PHDs which leads to the degradation of the HIF-1α protein. If PHDs are blocked, HIF-1α survives, moves into the nucleus, and acts as a transcription factor regulating hundreds of genes (22-23). Using a PHD inhibitor (PHDi) with a drug delivery system injected systemically into a non-regenerative mouse strain, we found that we could regulate HIF-1α levels and induce ear hole closure indistinguishable from that seen in the MRL mouse (21, 24). Blocking HIF-1α using *siHIF1a* completely blocked the regenerative response (21). This was also true in the case of another soft tissue target, enhanced drug-induced liver regeneration (25).

To realistically approach the clinical problem of restoring defects through anatomically faithful skeletal regeneration as needed for instance in craniofacial injuries, we would require anatomically accurate bone regeneration to support the re-growth of overlying soft tissues. In this study, we show that a hydrogel formulation (24) of the PHDi 1,4DPCA (26) leads to a clinically relevant case of complete bone regeneration as well as soft tissue restoration. Using a mouse periodontitis model (27-29) which employs an externally applied tooth ligature to induce bone degeneration through bacterial accumulation and inflammation, the effect of 1,4-DPCA was tested. Typically, studies of bone healing employ models with defined surgical defects that often don’t capture the complexity of real-world injuries and disease processes. In this case, however, bone loss is induced by host bacterial overgrowth and immune mediated damage which emulates the natural course of disease seen in clinical periodontitis and without any surgical intervention. After ligature removal, once periodontitis was established (day 10), injectable drug was administered to mice at a distal site and the mice were then followed.

In this study using Micro-CT analysis of bone damage and regeneration, we explored the effect of a 1,4-DPCA-containing formulation: DPCA-PEG (24) for over 220 days. We examined multiple timepoints to determine if full regeneration was achieved. After the ligature was removed and drug initiated, we examined mice in longitudinal studies and found that by day 20 after drug initiation, not only was full rapid recovery seen with normal alveolar bone architecture but also soft tissue regeneration and an increase in stem cell markers. In both the pulp of the teeth and the periodontal ligament (PDL), the primary connective tissue attachment of the tooth roots to the bone, we found increased cell proliferation, increased HIF-1α levels, bone-PDL interactions, vascularization, and tissue specific progenitor and stem cell markers. These included 1) scleraxis, a transcription factor expressed in tendon progenitor populations and mature tendon and which has also been found in PDL fibroblasts (30-33) and 2) neurofilament, a structural protein of mature nerve fibers, was seen in the pulp and also found in the PDL (34). The known pulp markers of progenitor cell populations, shown to be mesenchymal stem cells (MSC), alpha smooth muscle actin (aSMA) and CD44 (35-38), were increased after drug treatment. Furthermore, the possibility of tissue de-differentiation followed by re-differentiation into mature tissue, a hallmark of classic epimorphic regeneration with the expression of OCT3/4, Nanog, PAX7, SOX2, CD34 and seen in the ear regenerative response, was also noted (1-3,21).

In the broadest sense, bone is the biological “bioscaffold” which structurally supports the soft tissues of the body. The ability to regenerate bone through the up-regulation of HIF-1α by a systemically acting drug extends the range of possibilities of regenerative therapies.

## Results

### Ligature-induced bone degeneration

All C57BL/6 (B6) female mice in these studies had a Micro-CT scan performed 2 days prior to placement of a ligature (day minus 2) around the maxillary left 2^nd^ molar (Fig.1Aab, ref 27). Mice were re-scanned on day 5 (with ligature present) and again on day 10 (immediately after ligature removal). Image analysis showed significant bone degeneration within 5 days which became readily apparent by day 10 (Fig.1Ba-c). Mice tolerated the ligature well, displaying no obvious ill effects such as reduced chow ingestion. Graphed values of bone loss are seen in Fig.1C.

### Changes in bone regrowth in mice given DPCA-containing drug

At day 10, the ligature was removed and the mice were separated into two groups: 1) the control group which was treated no further; and 2) a group injected twice, on days 10 and 18 with DPCA-PEG (24). All mice were then imaged by Micro-CT on days 15, 21, and 30 (Fig.S1).

Three separate experiments were carried out, and the Micro-CT results were quantified. The key measure of alveolar bone loss and subsequent bone regeneration is derived by calculating the changes in bone area assessed from lateral images using the cementum-enamel junction (CEJ) and the crest of the buccal alveolar bone as anatomic landmarks (Fig.1D, E; Fig S1) to determine changes in each jaw. A statistical analysis was performed on the areas, and a statistically significant difference in the changes in bone area of the treated groups and the control group was seen. These differences clearly demonstrate the efficacy of the drug used in the treatment group versus the control group in reversing the induced periodontal disease in this mouse model. In Fig S1, the jaws are shown digitally overlaid and aligned on top of one another and one can observe the changes that occur. In Fig 1D, the treatments and degree of increase/decrease in area in mm^2^ is seen, where highly significant changes occur with drug treatment.

**Figure 1.**
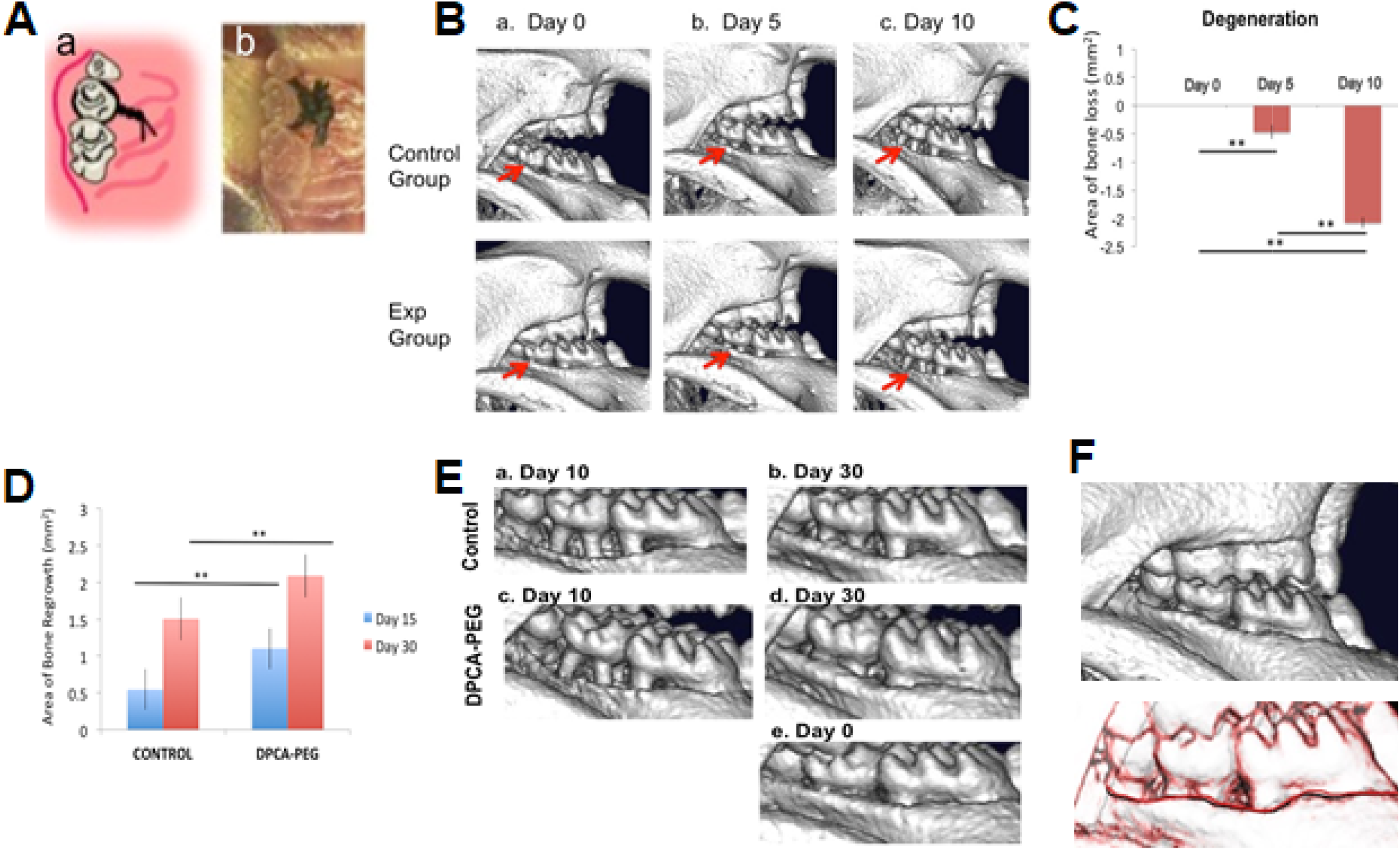
Degeneration of the mouse jawbone in the presence of ligature followed by regeneration of the jawbone post-ligature and post-drug. **Fig.1Aab**.The ligature-induced periodontitis model. 5–0 silk suture was passed through the interdentium between the first molar, the second molar and third molar using Dumont forceps. Suture was tied firmly using a triple-knot and excess suture was cut using spring scissors as seen in the cartoon (a) and photomicrograph (b). Taken from Ref. 27, Fig 1. **Fig.1Ba-c**. Micro-CT scans of jaws from mice during the 10 day ligature period for days 0, 5, and 10. The control group image is representative of the group (n=3) that will not receive drug (control, upper row) and the experimental group image is representative of the group (n=3) that will receive drug (experimental, lower row). The red arrows show the maxillary left second molars with bone degeneration extending to the adjacent 1^st^ and 3^rd^ molars. For visual clarity, the images are inverted 180 degrees (now: mandible top, maxilla bottom). **Fig.1C**. The area of degeneration was determined for all animals tested and shown for day 5 and day 10 after ligature placement. The y-axis is: Area of bone loss (mm^2^); (n= 7); error bars represent standard errors; for days 0-5, *p* = 0.00289; for days 0-10, *p* = 7.33084E-11; and for days 5-10, *p* = 3.60818E-07. (**) represents p<0.01. **In Fig 1D**, a graph of area of bone growth is seen for mice post–ligature but not given drug (no drug control) vs mice given drug (DPCA-PEG). Mice were injected with DPCA-PEG subcutaneously on days 0 and 8 after ligature placement and removal. MicroCT scans were obtained on day 15 of the experiment (day 5 post-ligature) and on day 30 (day 20 post-ligature). Here, a statistical analysis of the area of bone growth (mm^2^) is seen. Significant differences are found between the no drug control mice (n=10) and DPCA-PEG-treated mice (*p*=0.00253) on day 15 (blue bars) (n=12). The same is true on day 30, where DPCA-PEG-treated mice showed highly significant differences from non-drug-treated controls (p=0.00612) (red bars). Area analysis was performed as described in the Materials and Methods. The Y-axis = Area of bone regrowth (mm^2^); the error bars represent standard errors; and p values are represented as (*) = *p*<0.05; (**) = *p* <0.01. Mouse jaws analyzed were n=10 for ligature, no drug; n=12 for ligature/plus drug. **In Fig.1E**. micro-CT data shows a representative mouse maxilla which had ligature removed at day 10, scanned on day 10 and then re-scanned on day 30 (**Fig.1Ea,b**) as compared to a representative mouse maxilla receiving ligature and DPCA-PEG drug seen on day 10 and day 30 (**Fig.1Ecd**). The level of regrowth in the drug-treated group shows an almost, if not complete, return to what is seen before the start of the experiment (da0) (**Fig.1Ede**). **In Fig 1F**, there was no change in bone histology approximately 6 months later. Mice injected with DPCA-PEG drug were kept for additional observation as they aged. Over six months after the da 30 scan, mice were rescanned (**upper panel**) and then compared to the day 30 scan by overlaying the two scans on da 30 and da 220 (**lower panel**). The black line is the da 30 scan and the red line is the da 220 scan. Though shown as two lines, they are exactly overlapping. This result is representative of three mice.

From the above studies, DPCA-PEG showed rapid and significant bone regrowth. Comparing the jaw from the day that the ligature is removed where unusually extreme jawbone degeneration is seen by day 10 in the example shown in Fig.1Ea,c, one can see dramatic bone re-growth (Fig.1D,Ed, Day 20 after ligature removal). The regrowth of alveolar bone in mice given DPCA-PEG is nearly, if not completely, recovered when compared to the image taken before ligature and drug treatment, day 0 (Fig.1Ee). It is important to note that not only does the bone length and the apparent bulk volume return to normal within 30 days, but also the morphology of regenerated bone is indistinguishable from normal alveolar bone. Thus, the thickened boney alveolar ridge adjacent to the crowns is fully restored in drug-treated mice.

It should also be noted that the control bone itself grows back to some extent, consistent with earlier observations (39-40). This is not surprising since it has been long established that many rodent species continually wear down tooth crowns and there is subsequent regrowth of tooth roots and surrounding alveolar bone (see Discussion).

Finally, three of the DPCA-PEG-injected mice were examined for any long-term effects or reversal of the healed injury. After approximately six months from the end of the experimental time period (day 30), bone morphology was still stable. As seen in Fig.1F, no reversal of healing was seen and no adverse effects of the drug to the health of the mice were seen.

### Soft tissue changes post-ligature and post-drug

Bone loss in PD is preceded by bacterially-induced inflammatory changes in the soft tissue (gingiva) characterized by swelling and bleeding edematous tissue. As bone loss continues, the inflammatory state is intensified with breakdown of attachment fibers between the supporting alveolar bone and the roots of the teeth. The combination of direct bone loss and breakdown of periodontal (PDL) fibers leads to tooth mobility and eventual tooth loss. Thus, it is important to observe whether drug-induced regenerative therapy not only restores bone, but also restores soft tissue integrity.

In Fig.2Aa-c and Fig2Da, normal tissue histology around the tooth is shown with rete pegs in the epidermis, un-inflamed dermis, rich cellular pulp, and a PDL surrounding the full tooth root.

In Fig.2Ba-c, a ligature has been tied around the second molar for 10 days. At twenty days after ligature removal (day 30), there is significant ablation of the PDL root-bone attachment down to the root apex (green arrows). Furthermore, there is increased porosity of the surrounding alveolar bone (Fig 2Db). The human clinical correlate in this example would be extreme PD where tooth extraction would be indicated. Thus, compared to some alveolar bone recovery as seen by micro-CT (Fig.1Eab), we note that soft tissue recovery is poor by day 30 (Fig 2Ba,c; Fig 2E).

**Figure 2.**
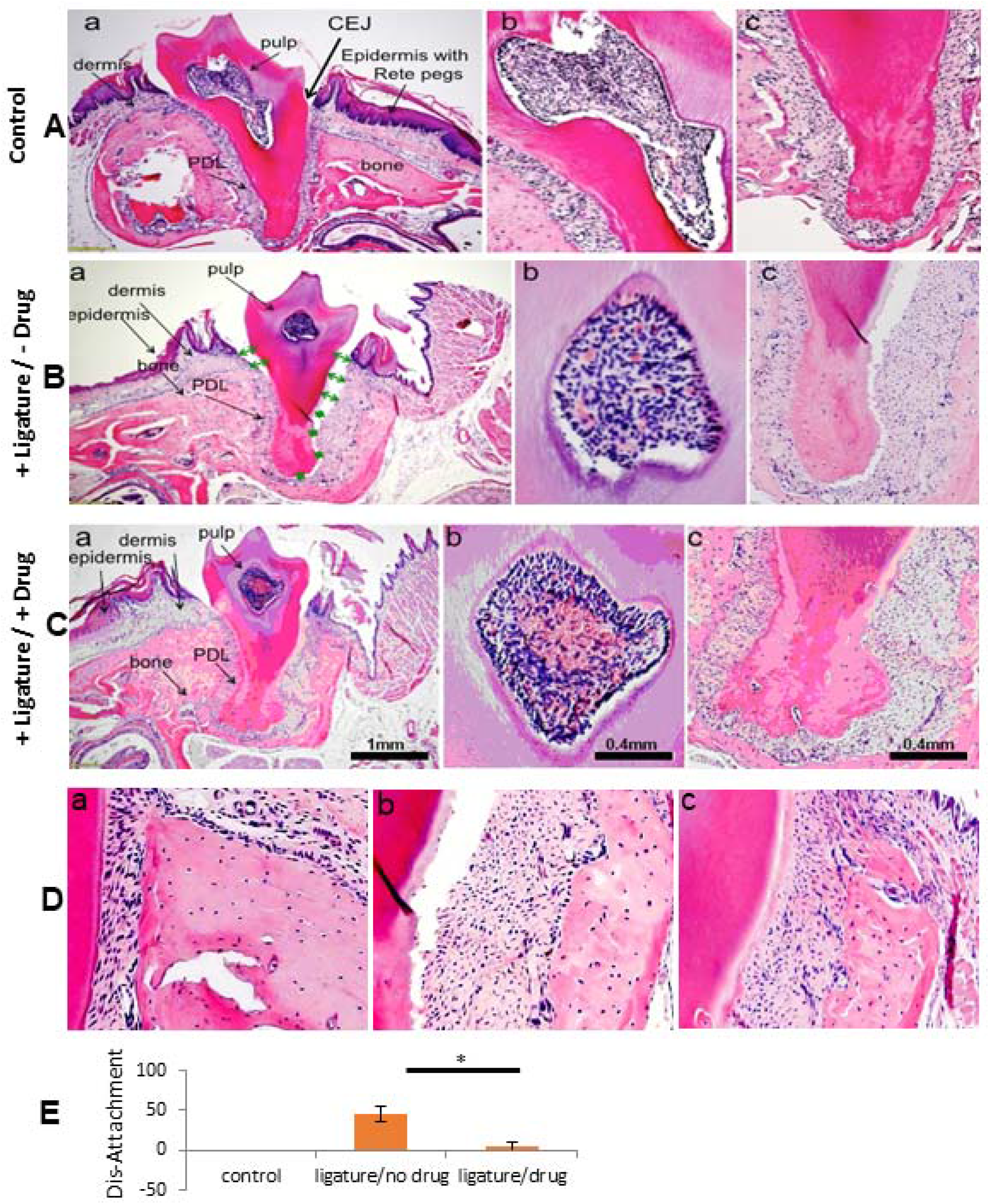
Soft tissue histological analysis of normal jaw and 20 days post-ligature +/-drug. In **Fig 2**, H&E-stained lateral jaw sections of a second maxillary molar in seen a 1) normal mouse maxilla (**Fig.2Aa-c)**, 2) maxilla from a +ligature/-drug-treated mouse (**Fig.2Ba-c)**, and 3) maxilla from a +ligature/+DPCA-PEG-treated mouse (**Fig.2Ca-c)** are seen on experimental day 30 (day 20 post ligature) and is representative of 3 mice. In the normal control mouse (**Fig.2Aa**) is seen a thick gingival dermis and epidermis with Rete pegs, a rich periodontal ligament attachment extending from root tip to cemento-enamel junction (CEJ), with periodontal ligament fibers and fibroblasts attached from tooth to normal bone (**Fig.2Da**), along with pulp chamber and its rich cellular composition, different from ligature-treated mice (**Fig.2Ba-2Ca**). The pathology induced from a silk ligature around the root adjacent to the crown is seen in **Fig.1Aab. Fig.2B and 2C**. In the +ligature-treated/-drug-treated mouse (**Fig.2Bac)**, a space between PDL and tooth extending to both sides of the tooth apex (green arrows, stars) is seen. Breakdown of dermis and epidermis surrounding the tooth crown with obvious dis-attachment of PDL to root surface results in a deep invagination extending > 60% of the length of the CEJ to tooth APEX. PDL is totally eliminated on the right of **Fig.2Bac** and partially obliterated on the left side. Epidermis and dermis are tattered and edematous. Bone shows porous changes (**Fig2Db**) and clinically this tooth would be highly mobile within the tooth socket and correspond in humans to advanced PD involvement within an indication for extraction. The jaw in **Fig.2Ca** is from a +ligature/+drug-treated (2 doses DPCA-PEG, da0, da8 post-ligature) mouse. Here, PDL is attached to bone (**Fig 2Dc**) and tooth, dermis is richer than without drug-treatment. Pulp shows higher levels of blood cells and vessels. A higher magnification of pulp shows differences with different treatments (**Fig.2A-Cb**). After drug treatment, the pulp is richer with higher levels of red cells and angiogenesis compared to both normal and ligature alone-treated mice (pink/red).

Fig.2Ca-c shows tissue from mice treated with ligature and then drug. On day 30, we now see a recovering cell-rich reattached PDL fully surrounding the root (Fig 2Cc, Fig2E), alveolar bone with less porosity (Fig 2Dc), and a highly vascularized pulp (Fig 2Cb). PDL fibers have returned (Fig 2Cc and 2Dc).

### Vascular changes in pulp and increases in stem cell markers post drug

Twenty days after drug initiation (day 30), there was an almost 8-fold increase in cellularity and vascularity in the B6 pulp after drug versus the no drug control (Fig.2Cb; Fig.3a). Neurofilament IHC, used to measure nerve in the highly innervated pulp, showed a significant increase in B6 mouse jaws given ligature and drug (Fig.3e). Also, increases in early stem cell markers such as OCT3/4 and PAX7 (Fig.3c,d) and the mesenchymal stem cell marker aSMA (Fig.3f) were also seen. Two markers, SOX2 and CD34 showed no differences with or without drug.

**Figure 3.**
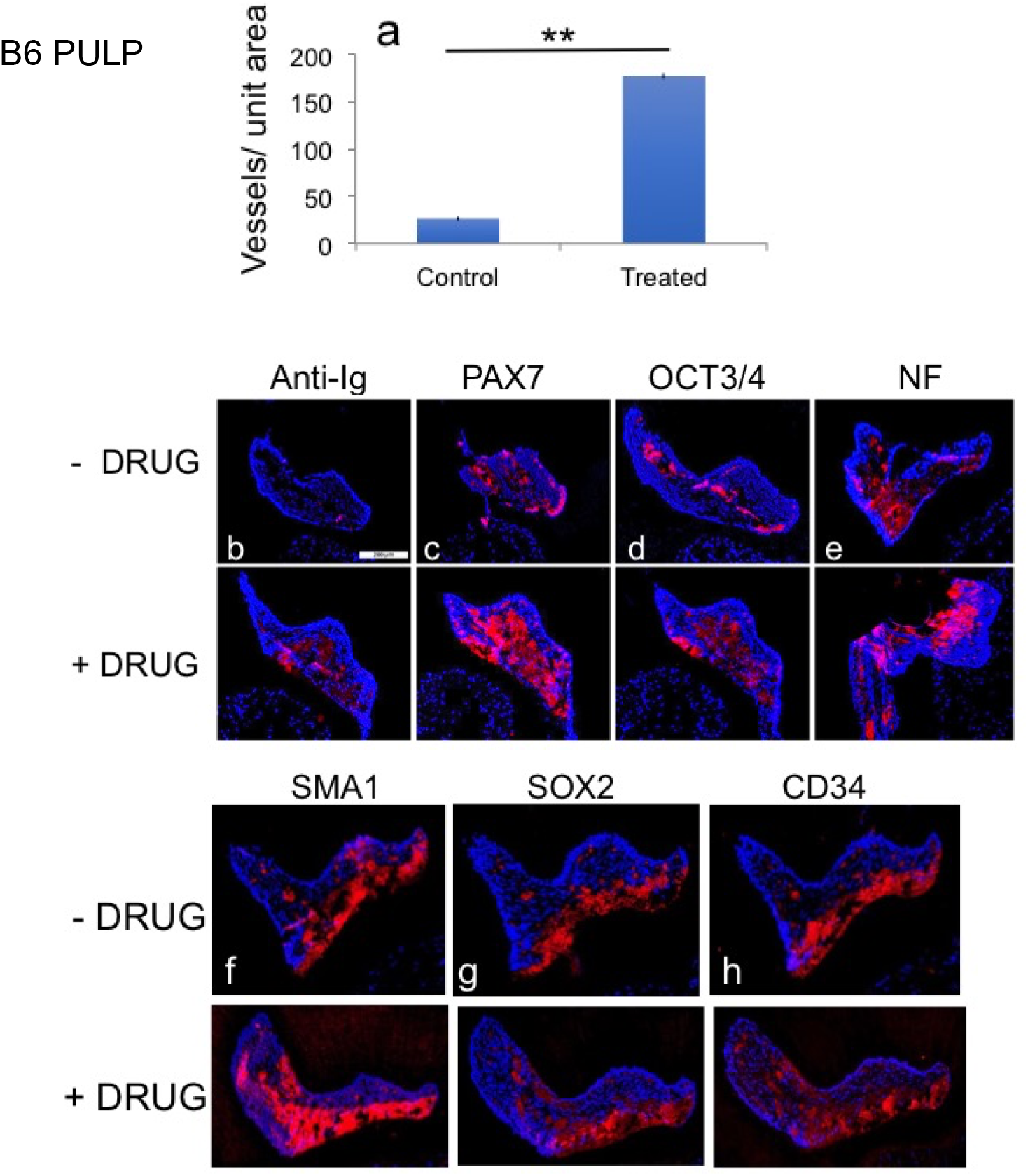
Marker expression in pulp after ligature and drug-treatment. In **Fig.3a**, a graph of the level of angiogenesis in the pulp (**Fig2Bb-2Cb**) is presented. The level of red in the pulp was determined by determining the number of pixels of red (IHC staining) using Photoshop as compared to the number of pulp area pixels (blue, DAPI) and is shown as a graph of the level of red in Fig.3a. The Y-axis is the vessel area/pulp area in the control and drug-treated mice; n=4; the error bars are standard errors; the *p*=0.0027127 and (**) represents p<0.01. Stem cell marker expression for PAX7, OCT3/4, aSMA (SMA1), SOX2, and CD34 was determined by IHC (red) in the pulp and is seen in **Fig.3b-h**. The anti-Ig control is lower in drug-untreated than treated samples. The scale bar = 200 um. The pulp was also examined for expression levels of neurofilament (NF), an intermediate nerve fiber filament and marker of innervation (**Fig.3e**). Quantitative staining using Photoshop CS6 determined the # red pixels over the total number of pixels in the pulp giving % positive staining; (n=4) and percentage of red pixels to total pixels determined. For **Fig.3b-h**, the no drug control versus plus drug experimental is seen for b (0.7% vs 6%); c (14% vs 38%); d (9% vs 21%); e (17.5% vs 27%); f (19% vs 38%); g (12.5% vs 13%); and h (18% vs 18%). Two markers, SOX2 and CD34, did not show a difference with and without drug at this timepoint.

### Changes in PDL markers

A closer examination of the B6 PDL showed that after ligature and drug treatment, the number of PDL fibroblasts increased two-fold compared to ligature alone. This was carried out examining longitudinal cross-sections (Fig.4a) and transverse cross-sections (Fig S2), both giving the same result

**Figure 4.**
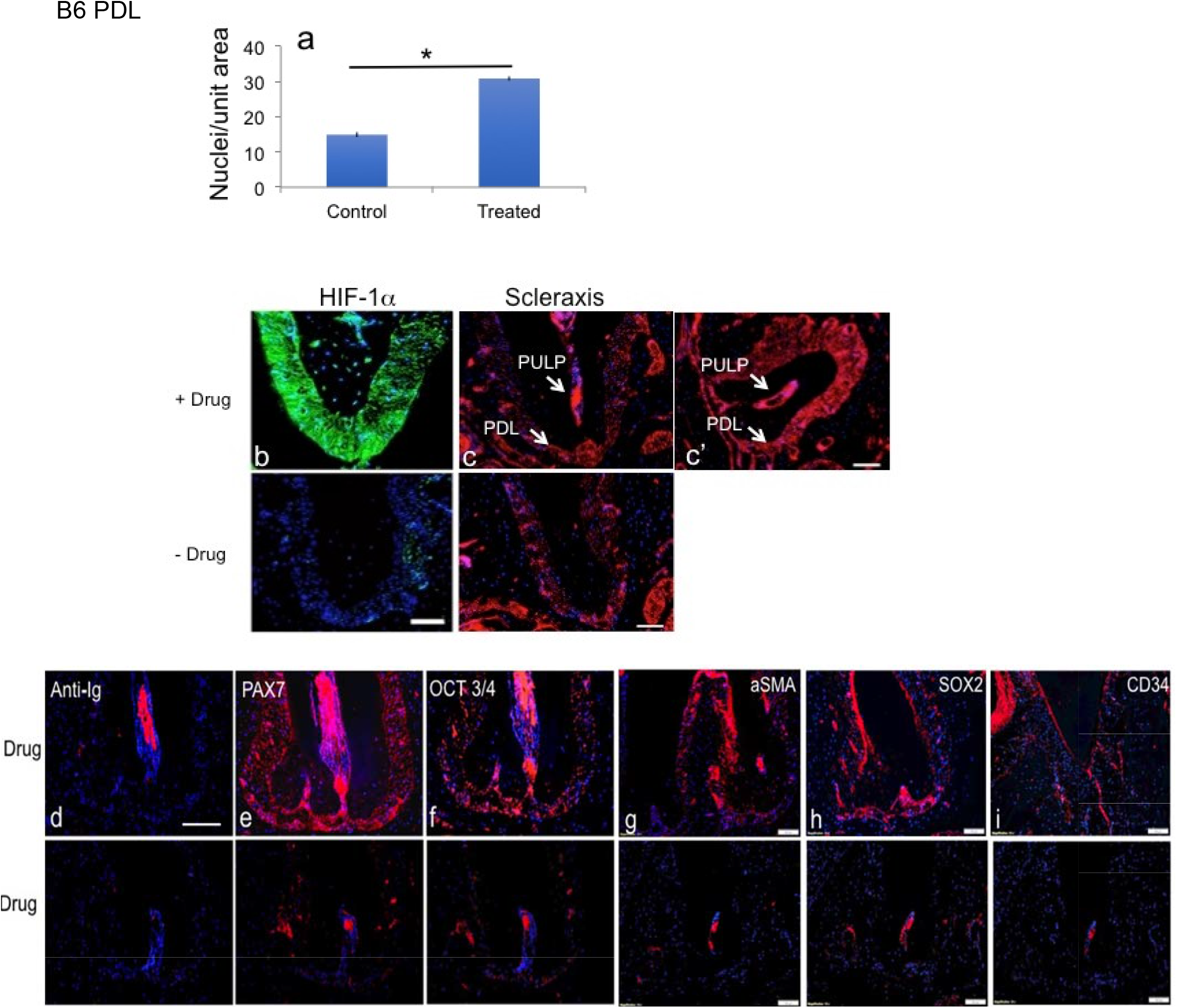
Marker expression in PDL after ligature and drug-treatment. In **Fig.4**, PDL analysis was carried out in B6-treated mice. In **Fig.4a**, in B6 mice, the number of PDL fibroblast nuclei were counted from **Fig.2BCc** and are two-fold greater in the drug-treated B6 mice. The Y-axis is the number of nuclei/unit area; error bars are standard errors; n=4; * *p*<0.01. In **Fig.4bcc’**, longitudinal jaw cross-sections from ligated-plus-drug (upper panels) versus ligated only (lower panels) respectively were stained with antibody to HIF-1α (green) and scleraxis (red) with PDL staining levels higher in sections from no-drug vs drug-treated mice (4% vs 11% red pixels; as described in **Fig.3**) (**Fig 4c**). The scale bar = 100um. In **Fig 4e-i**, antibodies to the stem cell markers PAX7, OCT3/4, aSMA, SOX2, and CD34 stained more highly in the drug-treated PDL (upper panels) compared to the non-drug treated PDL (lower panels). Anti-Ig controls showed no staining in the PDL except in the root canal (**Fig.4d**). The scale bar = 100um.

Given that 1,4-DPCA blocks PHDs and leads to HIF-1α stabilization, we noted high HIF-1αexpression levels in the PDL as well as scleraxis expression, a marker of activated PDL (Fig.4bcc’). Post ligature and drug treatment, high expression levels of scleraxis was also observed in the pulp chamber and canal. In previous studies (21,24), after drug treatment and a regenerative response, we noted expression of de-differentiation and immature cell markers. Here, we noted that in both the PDL and pulp including the apical pulp canal, all of which are known to contain stem cell progenitors, there was increased immunostaining by the stem cell markers PAX7 and OCT3/4 (Fig.4e,f), and the MSC markers aSMA, SOX2, and CD34 (Fig.4g-i)(though not seen in the pulp for SOX2 and CD34).

Interest in the cellular location of stem cell markers, ie the cytoplasm vs the nucleus, could not be determined using tissue sections. Therefore, In-vitro analysis of stem cell marker location in cell culture was examined pre and post addition of 1,4-DPCA (Fig 5). Using gingival cells in the absence of DPCA (Fig 5A,A’), we found OCT3/4 expression in the cytoplasm but not in the nucleus. On the other hand, after treatment with DPCA (100uM) in-vitro, all cells showed nuclear localization of OCT3/4 as well as in the cytoplasm (Fig 5B,B’). The amount of OCT3/4 staining increased after DPCA treatment (Fig5C). In addition, after DPCA treatment, the cells and nuclei became smaller and changed shape. (Fig 5D). This phenotype would be expected in stem-like multipotent cells.

**Fig. 5.**
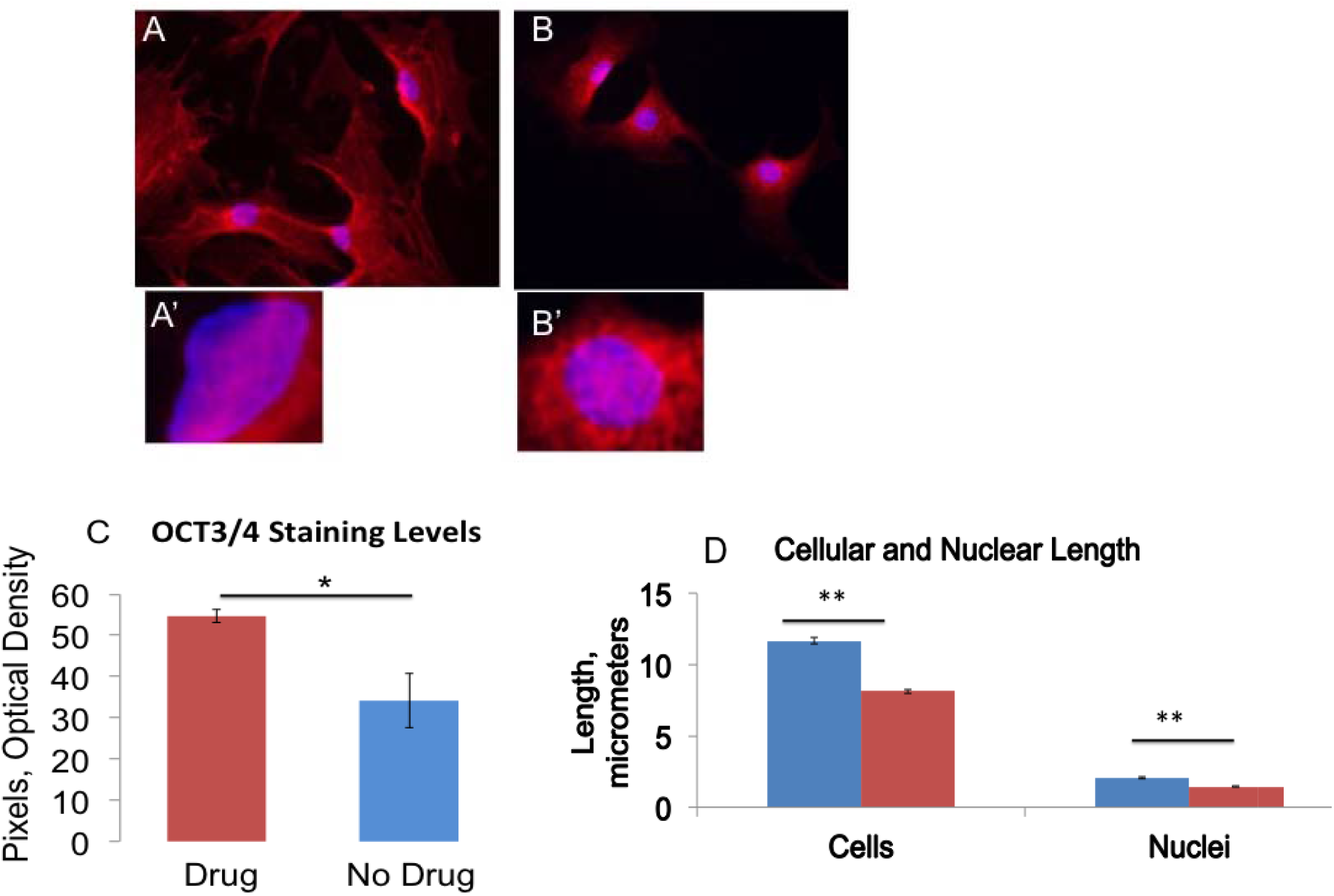
In-vitro nuclear expression of stem cell markers. In **Fig 5**, gingival cells from normal B6 female mice were grown on coverslips, treated without (A, or with 1,4-DPCA (100uM) (B, B’) overnight in culture, and stained with antibody to OCT3/4 (red) and the nuclei stained with DAPI (blue). The intensity of OCT3/4 staining for cells treated with drug vs cells untreated (normal) is seen in **Fig 5C**, p=0.0378 (n=3). The size of the cells and the nuclei (n=10 for each group) without and with DPCA treatment are shown in Fig 5D (blue bars=no DPCA; red bars = + DPCA), p=0.01 for change in cell length and p=0.003 for changes in nuclei length. (**) = p<0.01; (*) = p<0.05. Error bars = standard error.

## Discussion

### Induced periodontal disease as a model for bone and soft tissue regeneration

Bone, by its very nature is an exclusively internal organ system. Any laboratory intervention to create a model bone injury for wound healing studies, with the exclusion of blunt trauma injuries, necessarily requires surgical incisions through the overlaying skin, fascia and muscle. Periodontal disease, however, arises from a bacterial overgrowth of resident intra-oral micro-organisms external to soft tissues (the gums) of the oral cavity with a subsequent host inflammatory response leading to massive lesions of the underlying alveolar bone. An induced periodontal disease model, is therefore attractive from the perspectives of both basic bone regeneration biology and as a translational stepping stone in the treatment of a common human disease.

With the declining incidence of caries, periodontal disease (PD) has emerged as the most common cause of tooth loss affecting 30-60% of the adult population, presenting a major clinical challenge (41,42) with the loss of supporting alveolar bone around the roots of teeth as well as the destruction of adjacent soft tissues (43). This can be localized to a single tooth or involve virtually the entire dentition. The etiology of PD is due to a number of identified periodontopathic organisms found beneath the gingival tissue (27-29,43) combined with unidentified or identified host susceptibility factors such as diabetes (44-45).

The challenge in the treatment of advanced PD, where bone loss may exceed 50% or even 80% of the length of the tooth root, is to halt further progression and achieve both a functionally and an esthetically acceptable outcome. Thus, a primary goal is to restore lost alveolar bone and soft tissues to their original state without the loss of teeth (46-47). It is for this reason that we have explored PHDi-based regenerative drug-induced bone and soft tissue regeneration in a PD model.

### The role of HIF-1αand changes in vascularization

The central molecular regeneration pathway as understood from previous soft-tissue studies (21,24) involved in the use of PHDi is the up-regulation of HIF-1α. HIF-1*α* target molecules include *Pdk*, and *Ldha, Pkm2*, and *Gapdh*, all glycolytic enzymes, and leading metabolic remodeling (22-23). HIF-1*α* was identified as the molecule responsible for the broad regenerative ability of the MRL mouse through multiple findings. In genetic mapping studies of MRL mice, a reduced level of the gene RNF7, a part of the ubiquitin ligase complex, and necessary for proteolysis and lowered protein expression levels of HIF-1α (48), proved to be a candidate gene associated with regenerative healing (49). Furthermore, MRL mice were found to be metabolically more embryonic using aerobic glycolysis (19,20). Finally, *siHIF1a* completely suppressed ear hole closure both in the MRL and non-regenerative mice treated with drug (21).

One of the many intriguing aspects of HIF-1α elevation is the de-differentiation of mature cell populations to an immature state determined by molecular markers and aerobic glycolysis (50-57). This state could be key to the regenerative response. Also, HIF-1α is responsible for an enhanced vascularization response, producing molecules such as VEGF and HMOX1. It is clear in the pulp that vascular tissue is significantly increased 15-20 days after drug administration (Fig 2Cb, Fig 3a). This change in vascularity could lead to an increased number of stem cells migrating into the tooth as noted by stem cell marker increase (Fig 3b-f). On the other hand, DPCA-PEG could lead to de-differentiation with increased stem cell markers and growth as seen in the ear accompanying regenereative ear hole closure (21, 24).

### Induction of stem cell markers

Many molecular and cellular markers of regeneration observed in spontaneously regenerating species such as the newt and axolotl (1-3) and in the MRL mouse (19) were indistinguishable from those seen in drug-treated mice (21,24). Our previous studies largely focused on soft tissue, specifically in the ear pinna and included the regrowth of hair follicles and cartilage (21,24). In the PD model studied here, soft tissue is also affected by the drug, and multiple early impacts of the drug 5 days after ligature is removed included increased Treg FOXP3+ populations and lowered inflammatory cytokines (58). In the current study, 20 days after ligature is removed, the PDL, which is key to supporting the teeth and securing them to the bone, after drug treatment showed complete re-attachment to the teeth with increased numbers of PDL. On the other hand, ligature without drug leads to PDL separated from the teeth without reattachment, typical of human PD. The PDL after drug administration also shows increased levels of scleraxis, a transcription factor considered to be a marker of PDL (30-33), a marker also seen associated with osteocytes and cementoblasts (32).

Another dental soft tissue target, the pulp, is considered to be a source of stem cells and contains dentinoblasts which “respackle” the inside of the tooth dentin. We noted that the pulp vasculature was increased after drug as well as stem cell markers. As mentioned above, whether this is due to a de-differentiation process in the pulp or to increased numbers of stem cells in the over-abundant vasculature is not clear. We also noted an increase of neurofilament (*34*), a nerve marker, which is present in normal pulp and increased after drug.

Stem cell populations have been identified that are associated with teeth in the pulp chamber, at the base of growing roots, and in the PDL (59). Progenitor cell populations have been identified in periodontal tissue previously and express mesenchymal stem cell markers such as STRO-1, CD146, CD44, and αSMA (30-33,60-62). These progenitors exhibit many stem-cell-like features, including small size, responsiveness to stimulating factors, slow cycle time and the ability to generate multiple mesenchymal lineages (63-65). In addition, neural crest-derived cells have been identified in the periodontal ligament and the pulp chamber using markers such as Slug, AP2 alpha, HNK-1, p75NTR and Nestin (66-70).

Alpha SMA has been found in stem cells and regenerating tissue as well as in blood vessel pericytes and myoepithelial cells involving force-generating function. During mandible development, aSMA was found in the dental follicle and then in the periostin-positive area along with RUNX2 positive cells and localize on the alveolar bone region suggesting involvement in bone formation (71).

We tested multiple stem cell markers such as OCT3/4, PAX7, SOX2, CD34 and aSMA finding the same de-differentiation markers in both pulp and PDL and these markers increased after giving the drug. These markers could be due to the dedifferentiation of mature cells (1-3,21,24) or could be markers of stem cell populations in the pulp and PDL as discussed above (72). Cross talk between the pulp and the periodontal ligament should not be ruled out since the pulp and PDL are anatomically connected via the apical root vasculature.

Although stem cell markers were increased in the PDL after systemic delivery of 1,4-DPCA-PEG, it was difficult to see cellular sub-structure and intra-cellular location of these stem cell markers. Therefore, we grew gingival-derived fibroblast-like cells to analyze the staining of OCT 3/4 with and without addition of DPCA. Here, we found that the cells before DPCA showed staining in the cytoplasm but after DPCA, staining was seen both in the cytoplasm and in the nucleus where Oct 3/4 acts as a transcription factor and is key to self-renewal of undifferentiated embryonic stem cells and is a specific marker for dedifferentiation (51). One interesting effect of DPCA on the gingival cells in-vitro is that the cells go from a culture of large multinucleated cells to a culture of small mononuclear cells. This is reminiscent of a classical regeneration model in-vitro in which rat multinucleated and terminally-differentiated skeletal myotubes separate into small mono-nucleated cells after treatment with an amphibian regeneration blastema factor or msx-1 (73-74).

Taken together, our present work has shown the ability of systemic DPCA-PEG therapy to completely regenerate severely degraded alveolar bone and soft tissue with remarkable anatomic fidelity in 20 days.

It should be noted that in humans, regenerative recovery of bone is seen, but is usually limited to bone fractures which are aligned with a gap of less than 1 mm (75-76). On the other hand, it is also known that both the maxillary and mandibular incisor teeth of rodents kept in animal colonies on soft chow often elongate (33,34) and we are seeing a background growth effect. In studies reported here however, the regenerative contribution of the drug is clearly distinguishable from background.

In conclusion, this study extends our previous work on drug-mediated stabilization of HIF-1α to achieve soft tissue regeneration in mice (77) in new directions. First, 1,4-DPCA delivered systemically rapidly reverses severe bone loss in an anatomically complex structure, the alveolar bone of the maxilla, leading to a perfect replica of healthy bone and associated soft tissue such as PDL with long-term maintenance. Second, the induced bone loss occurs in an experimental system which emulates the bacterial etiology of the human disease, periodontitis.

## Materials and Methods

### Study Design

We used the inbred mouse strain, C57BL/6 female, to study the effect of a small-molecule inhibitor of PHDs on the in-vivo expression of HIF-1α and the impact on quantitative regenerative maxillary bone growth *(58)* and soft tissue regeneration including PDL and pulp. A 5.0 silk ligature model was used to induce periodontitis in the mice and DPCA–PEG was tested and subcutaneously implanted in the back of the neck at multiple time points (Fig.6). End points of the study were previously determined to be a minimal of 30 days and up to 50 days after injury and included key indices of tissue regeneration such as bone regrowth as measured by Micro-CT scanning, soft tissue regrowth after H&E histological analysis including epidermis, gingiva, PDL attachment and number, and markers of regeneration determined by immunohistochemistry of jaw tissue for stem cell markers, and HIF-1α, neurofilament, and scleraxis and RT-PCR analysis of gingiva. These parameters involved physical measurements of growth and gene expression at the RNA and protein levels. Tissue was coded and different laboratory personnel were involved in doing ligatures, scanning, tissue preparation, and data analysis.

**Figure 6.**
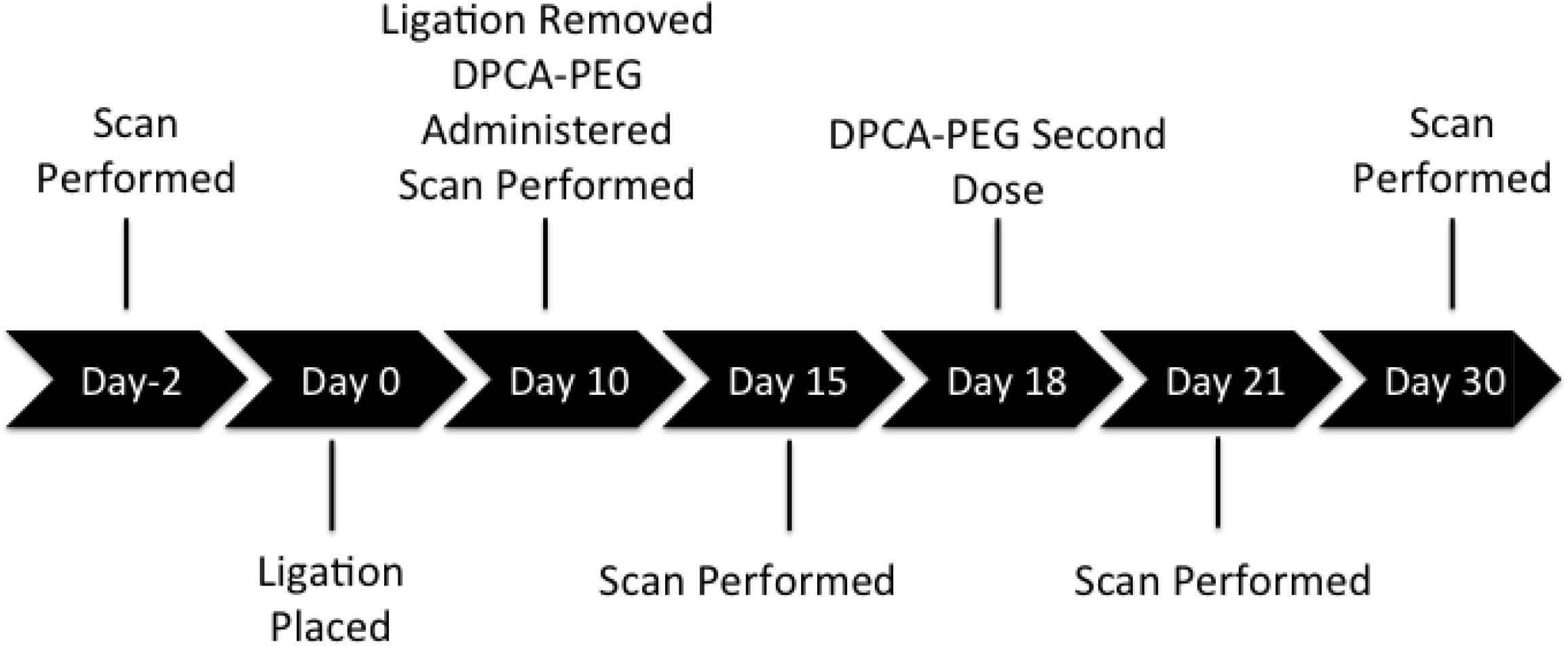
Timeline of treatments.

### Animals

C57BL/6 female mice, 9 weeks of age, were obtained from Taconic Laboratories. The experiment was done with 2 groups of 3-4 mice based on the previous work of Hajishengallis et al. (27) and was repeated twice for a total of 10 mice in each group. A previously described drug construct (24) was tested for its ability to regenerate periodontal bone, a recent polymeric 1,4-DPCA construct with improved rheology (24) compared to our original drug construct (21).

Food and water were provided ad libitum. All animal experiments were reviewed and approved by the Institutional Animal Care and Use Committee of LIMR and were performed in compliance with institutional, state, and federal policies.

### Tissue Culture

Primary fibroblast-like cell lines from gingival tissue were established from B6 female mice by plating in dispase and then collagenase and then grown in DMEM-10% FBS supplemented with 2 mM L-glutamine 100 IU/mL penicillin streptomycin and maintained at 37 °C, 5% CO2, and 21% O2. The cells were wash and only adherent fibroblasts maintained. Cells were split 1:5 as needed to maintain exponential growth avoid contact inhibition. Passage numbers were documented and cells from early passages (<P20) froz in liquid nitrogen and used in the described experiments.

For immunohistochemical staining, primary gingival fibroblasts were grown on coverslips in DMEM with 10% FBS at 37 °C in a humidified 5% CO2 incubator. Before staining, cells were incubated with DPCA overnight. The coverslips were rinsed with 1× PBS, the cells were fixed in cold methanol (−20 °C) for 10 min, rinsed with 1× PBS, treated with 0.1% Triton-X100, and then incubated with the appropriate primary and secondary antibodies (Table S1). Photomicrographs were produced using the fluorescent microscope (Olympus AX70) and a DP74 camera with cellSens Standard software for image analysis.

### Drug application

DPCA-PEG was synthesized as described (24) and was injected twice subcutaneously every 8 days (day 0 and day 8) using 25ul of gel #10/injection or about 50ug of DPCA. The time course of injections and scans performed during longitudinal experiments can be seen in Fig.6.

### Preparation and immunohistochemistry of jaw tissue

Tissue from upper jaws was fixed in Prefer fixative (the active ingredient is glyoxal) (Anatech) for 5 days and then washed in H_2_0. Jaw tissue was then decalcified using UltraPure 0.5 M EDTA, pH 8.0 for 5 weeks with changes in EDTA solution twice a week. Tissue was put in ETOH and then embedded in paraffin and 5-μm thick sections cut. Before staining, slides were dewaxed in xylene and rehydrated. Tissue sections were then treated with 3% H_2_O_2_ and nonspecific binding was blocked with 4% BSA (A7906; Sigma) for 1 h. The primary antibodies and matched secondary antibodies used for IHC were shown in Supplement A Table S1. Photomicrographs were produced using the fluorescent microscope (Olympus AX70) with an Olympus D74 Color Camera using CellSens imaging software.

For histological stains, tissue sections were treated as above and stained with Hematoxylin (Leica Microsystems, #3801562) and Eosin (Leica Microsystems, #3801602). The slides were washed, rehydrated, cleared with Xylene and coverslipped with Permount mounting media (Fisher, SP15-500). Staining was visualized using an Olympus (AX70) microscope in bright field for H&E and fluorescence.

### Induction of periodontal disease

For ligature placement, mice were anesthetized with Ketamine/Xylazine, a 5-0 silk ligature was placed around the upper left second molar of the maxilla according to an established procedure (27,58) generating a dental plaque-retentive milieu that reliably and quantitatively produces a periodontal bone lesion. Importantly, this ligature procedure induces not only bacterial over-growth but also selective expansion of periodontopathic microorganisms, mimicking quantitative and qualitative (dysbiotic) microbiome alterations, the same etiology as that in human disease (27,58). Upon placement of the ligature, the 5-0 silk ligation accumulates dental plaque and oral bacteria creating a local inflammatory state in the surrounding gums and bone of the tooth. Ketamine/Xylazine was used to remove the ligature and mice were re-scanned. Mice were subsequently scanned for further analysis. During this time, mice were kept on a normal diet of mouse chow and were monitored daily for any signs of physical discomfort in accordance with the Lankenau Institute for Medical Research (LIMR) Animal Care Policies and Procedures Manual. At each time point, animals were imaged and analyzed with the area analysis described below. On four occasions the ligature model was not completed due to the 5-0 silk ligation falling off of the tooth. These mice were excluded from the study and the remaining mice were randomly distributed before any hydrogel injections.

### Ketamine/Xylazine Mixture

Stock solutions of Ketamine (100 mg/ml, Hanna) and Xylazine (20 mg/ml, Hanna) are prepared using PBS as a diluent, respectively at a ratio of 3:1:16, and is vortexed and used immediately.

### MicroCT Scanning

For scanning, mice were anesthetized using isoflurane (Henry Schein) 2-4% concentration in 100 % O_2_ for 3 minutes in an anesthesia chamber. Upon establishing anesthesia, mice were placed into the microCT FX (Perkin Elmer; ref 78) tray with the isoflurane nose cone. 3D images were collected and rendered with a voltage of 90kV a CT current of 160 μA and a live current of 80 μA for 17 seconds time for a total dose of 11milliGY (78-79). This constitutes the pre-intervention baseline.

Buccal images were analyzed unlike original experiments done by measuring the palatal side without Micro-CT (27,57). They were normalized to the Micro-CT HA D4.5 phantom from QRM (Quality Assurance in Radiology and Medicine GmbH).

### MicroCT Analysis

An analytical method to quantitate longitudinal changes in bone morphology was designed and implemented. 3D renderings were obtained using Quantum software (Perkin Elmer) and subsequently converted to 2D images, which were then superimposed in Photoshop (CC 2019) for quantitation of bone morphological changes.

### Statistical analysis

For multiple-group comparisons, data were analyzed by one-way ANOVA followed by Tukey’s multiple comparison test. A two-tailed Student’s *t*-test was used for two-group comparisons. *P* values <0.05 were considered to be statistically significant and values <0.01 were considered highly significant. All statistical analyses were performed using RStudio (version 1.1.463) with the stats and stats4 packages (versions 3.5.1).

## Abbreviations

B6: C57BL/6 mouse
CEJ: cemento-enamel junction
DPCA: 1,4-dihydrophenonthrolin-4-one-3-carboxylic acid
HIF-1α: hypoxia-inducible factor-1alpha
MRL: Murphy Roths Large mouse
PD: periodontal disease
PDL: periodontal ligament
PHDi: prolyl hydroxylase inhibitor

## Author Contributions

EZ, TK and EHK designed, performed experiments, analyzed, interpreted, and graphed data, and prepared the manuscript. GH provided expert advice and experimental design, and together with PBM interpreted the data, and critically reviewed the manuscript. PBM and JC provided drug development and experimental data, and constructs; YZ and KB carried out all Swiss Webster studies, AA carried out all Micro-CT scanning; SB provided animal care and tissue preparation; and KB carried out histology and immunohistochemistry studies. DS provided clinical dental expertise and manuscript preparation. All authors have read and approved the manuscript.

## Supplement

**Table S1:** Primary and Secondary Antibodies Used in Immunohistochemistry.

|  | 1st antibody |  |  | 2nd antibody |  |  |  |
| --- | --- | --- | --- | --- | --- | --- | --- |
|  | Company | Cat. no. | Dilution | All from Molecular<br>Probe | Company | Cat. no. | Dilution |
| HIF-1α | Abcam | ab2185 | 1:1000 | Alexa Fluor 488 goat<br>anti-rabbit IgG | Molecular<br>Probe | A11008 | 1:200 |
| Nanog | Calbiochem | SC1000 | 1:150 | Alexa Fluor 568 goat<br>anti-rabbit IgG | Molecular<br>Probe | A11036 | 1:400 |
| OCT-3/4 | Santa Cruz | sc-5279 | 1:150 | Alexa Fluor 568<br>rabbit anti-mouse<br>IgG | Molecular<br>Probe | A11061 | 1:400 |
| CD34 | Bioss | bs-0646R | 1:200 | Alexa Fluor 568 goat<br>anti-rabbit IgG | Molecular<br>Probe | A11036 | 1:300 |
| Alpha<br>Smooth<br>Muscle Actin | Thermo-Fisher | PA1-37024 | 1:100 | Alexa Fluor 568<br>goat anti-rabbit IgG | Molecular<br>Probe | A11036 | 1:300 |
| PAX7 | R&D<br>Systems | MAB1675 | 1:50 | Alexa Fluor 568<br>rabbit anti-mouse<br>IgG | Molecular<br>Probe | A11061 | 1:400 |
| Neurofilament | Sigma | MAB200 | 1:50 | Alexa Fluor 594 goat<br>anti-mouse IgG | Molecular<br>Probe | A11005 | 1:200 |
| Scleraxis | Santa Cruz | A-7 | 1:50 | Alexa Fluor 568 goat<br>anti-mouse IgG | Molecular<br>Probe | A11036 | 1:1000 |
| Ki67 | Sigma | C8791 | 1:200 | Alexa Fluor 488 goat<br>anti-mouse IgG | Molecular<br>Probe | A10680 | 1:100 |
| CD44 | PharMingen | 01221D | 1:100 | Alexa Fluor 594<br>goat anti-rat IgG | Molecular<br>Probe | A11007 | 1:200 |
| SOX2 | Chemicon | AB5603 | 1:500 | Alexa Fluor 568 goat<br>anti-rabbit IgG | Molecular<br>Probe | A11036 | 1:300 |

**Figure S1.**
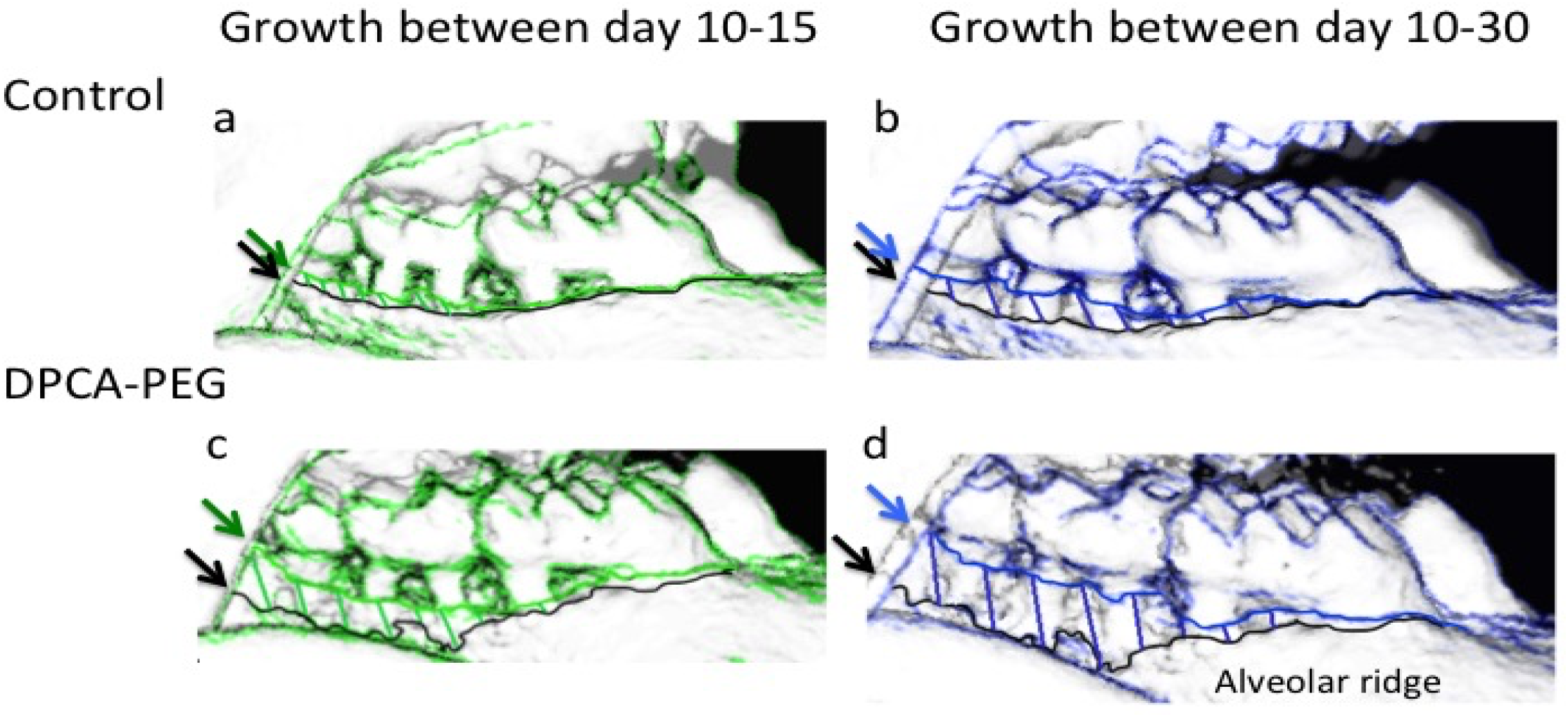
Overlays of jaws at different time points post-ligature. Using our analysis program, the changes in bone using Micro-CT scans and analysis appear as changes in bone using scans and overlays of jaws at day 0, 10, 15, and 30 after drug treatment. Micro-CT scans of jaws from control mice un-treated with DPCA-PEG and treated with DPCA-PEG (*18*) mice, 10 days after start of the experiment when the ligature is removed, are seen as black lines indicating bone levels at day 10 (black arrows), the green lines show bone levels determined by Micro-CT scans at day 15 (green arrows), and the blue lines indicate bone levels determined by Micro-CT scans at day 30 (blue arrows). A representative of the control group (n=3) which had the ligature on from day 0 to day 10 but no drug treatment, is seen in **Fig S1ab**. In (**FigS1c**) is the degree of growth between days10-15 and in (**Fig S1d**) is the amount of growth between days 10-30, seen in the drug-treated group (n=3).

**Figure S2.**
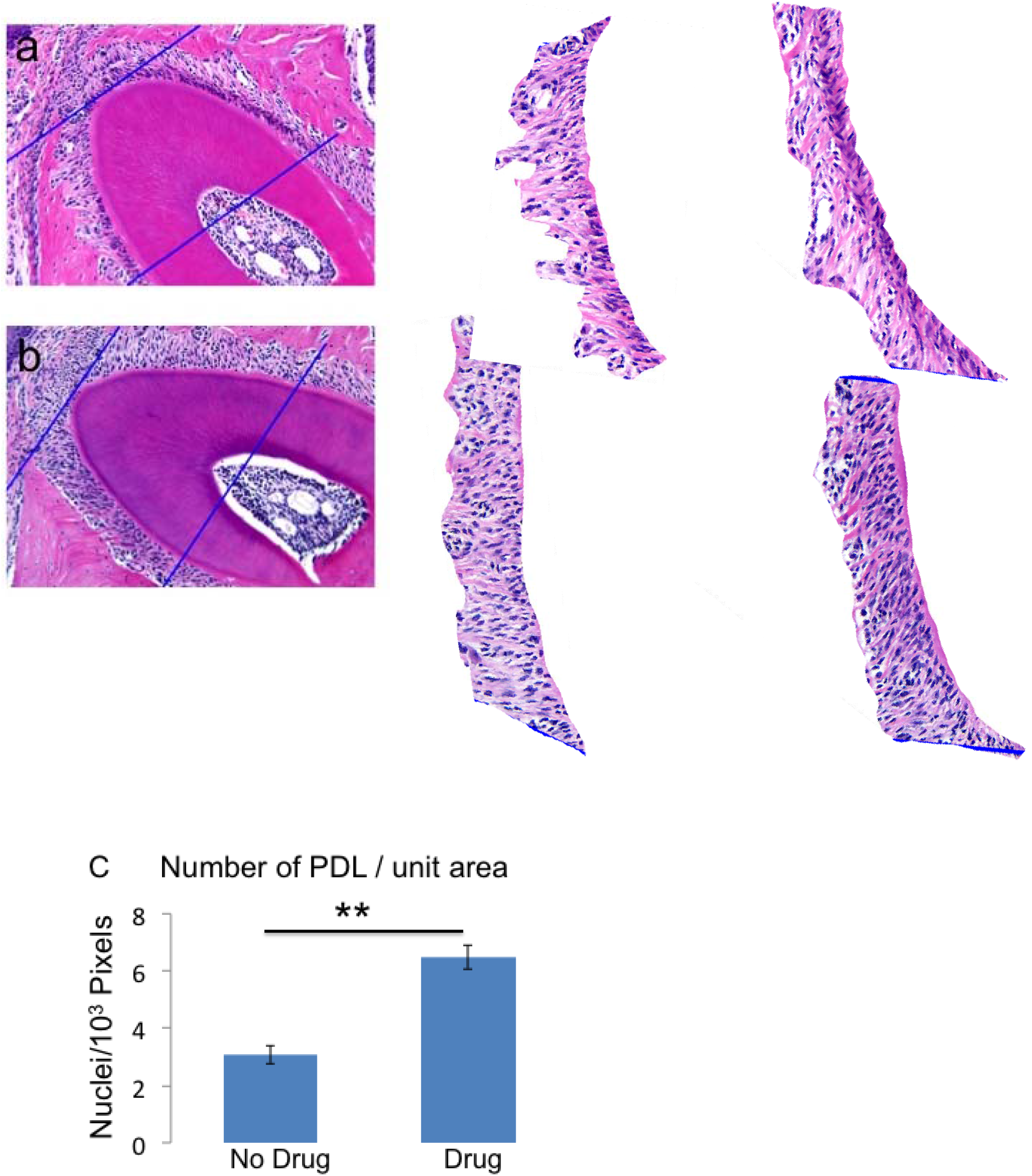
Counting PDL around the Root of the Second Molar. A transverse cross-section of the root of the upper 2^nd^ molar is examined and the same approximate area of the PDL is analyzed (**Fig S2a,b**). In **Fig.S2a**, PDL from mice without drug treatment is seen; in **Fig.S2b**, PDL from drug-treated mice is shown. The PDL area was cut out, enlarged, and fibroblast like-cell nuclei counted /unit area (**Fig S2c**) for ligature alone controls vs ligature plus drug-treated mice (n=6). p=8.44E-05 with (**) for p<0.01. Error bars = standard errors.

